# Allele-level visualization of transcription and chromatin by high-throughput imaging

**DOI:** 10.1101/2024.02.19.580973

**Authors:** Faisal Almansour, Adib Keikhosravi, Gianluca Pegoraro, Tom Misteli

## Abstract

The spatial arrangement of the genome within the nucleus is a pivotal aspect of cellular organization and function with implications for gene expression and regulation. While all genome organization features, such as loops, domains, and radial positioning, are non-random, they are characterized by a high degree of single-cell variability. Imaging approaches are ideally suited to visualize, measure, and study single-cell heterogeneity in genome organization. Here, we describe two methods for the detection of DNA and RNA of individual gene alleles by fluorescence in situ hybridization (FISH) in a high-throughput format. We have optimized combined DNA/RNA FISH approaches either using simultaneous or sequential detection. These optimized DNA and RNA FISH protocols, implemented in a 384-well plate format alongside automated image and data analysis, enable accurate detection of chromatin loci and their gene expression status across a large cell population with allele-level resolution. We successfully visualized *MYC* and *EGFR* DNA and RNA in multiple cell types, and we determined the radial position of active and inactive *MYC* and *EGFR* alleles. These optimized DNA/RNA detection approaches are versatile and sensitive tools for mapping of chromatin features and gene activity at the single-allele level and at high throughput.

## Introduction

Eukaryotic genomes are organized at multiple levels (Misteli 2020). The hierarchical organization, from nucleosomes to chromosomes, is thought to contribute to gene expression regulation through various features of chromatin organization, such as loops, domains, and preferred nuclear positions of gene loci (Gibcus and Dekker 2013; Bickmore 2013; Misteli 2020). The spatial arrangement of genomes changes dynamically during cellular differentiation, transcriptional activity, and disease progression (Shachar et al. 2015; Spielmann et al. 2018; Scholz et al. 2019; Finn and Misteli 2022), pointing to the possibility of a functional role of genome organization in gene regulation.

Fluorescence in situ hybridization (FISH) to detect specific regions of genomic DNA is an essential tool that has been widely used to probe the complex organization and 3D positioning of genes and chromosomes within the nucleus (Shachar et al. 2015; Finn et al. 2019). Complementarily, FISH detection of mRNA allows visualization of the transcriptional activity of individual gene alleles (Young et al. 2020). The combined application of DNA and RNA FISH allows for the concurrent assessment of chromatin features, for example, chromatin compaction or radial nuclear position and gene expression status. Detection of DNA and RNA at individual alleles is an important tool to address the key question of how genome organization relates to gene expression.

While some combined DNA/RNA FISH methods have been described (Lai et al. 2013; Barakat and Gribnau 2014; Petropoulos et al. 2016; Jowhar et al. 2018), they remain technically challenging because hybridization conditions for DNA and RNA detection are significantly different, typically necessitating sequential detection steps. In addition, to effectively cross-compare chromatin features and gene activity, a large number of individual alleles need to be measured to achieve high statistical power of analysis, requiring high-throughput imaging approaches. To bridge this gap, we have developed and optimized two high-throughput DNA/RNA FISH (DNA/RNA HiFISH) protocols using either a simultaneous or a sequential approach for visualizing DNA and RNA at the single-allele level (Fig. 1). These approaches allow the probing of the behavior of individual gene alleles, regardless of their activity status and enable the comparison of active and inactive alleles in the same cell nucleus and with high statistical power.

**Fig. 1.**
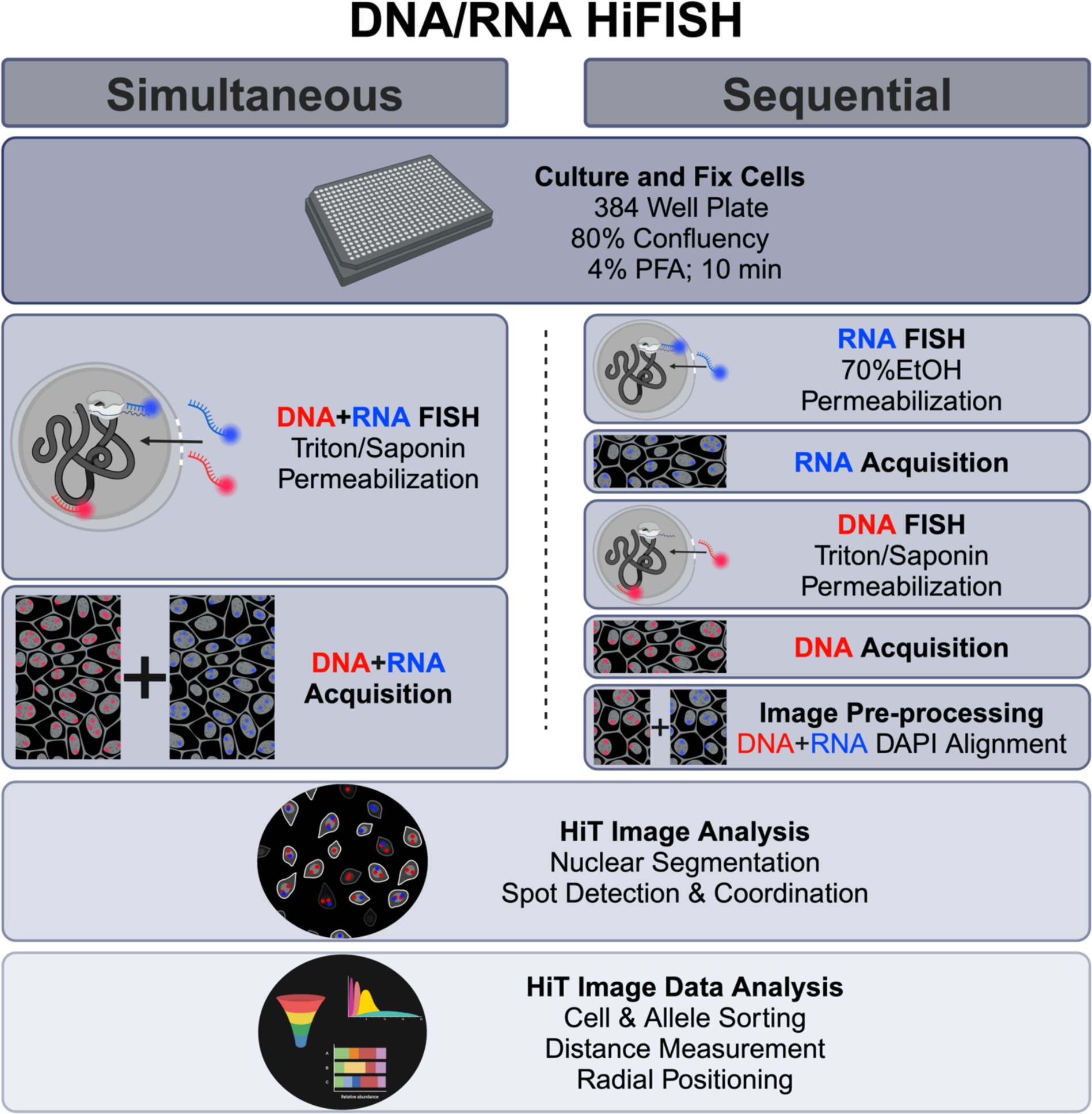
Outline of simultaneous and sequential DNA/RNA HiFISH approaches. The two approaches use the same cell culture and fixation steps, image processing, and image analysis but differ in the hybridization and image acquisition steps

Our protocols combine bacterial artificial chromosome (BAC) probes for DNA FISH probes with Stellaris® intronic RNA FISH probes against nascent mRNA in a 384-well plate format. In addition, we leverage automated 3D confocal microscopy and customized high-content image analysis workflows to quantitatively detect chromatin loci and gene expression status of individual alleles to distinguish active from inactive alleles. As proof of principle, we apply these protocols to high-throughput DNA/RNA HiFISH to detect *MYC* and *EGFR* genes in two human cell lines. We demonstrate high sensitivity of DNA and RNA detection, and we compare the radial location of active and inactive *MYC* and *EGFR* alleles. These approaches represent optimized tools for precise mapping of chromatin organization and gene activity at the single-cell and single-allele levels.

## Materials and Methods

### Cell culture

Human bronchial epithelial cells (HBEC3-KT), derived from normal human bronchial tissue, were immortalized through the stable introduction of expression vectors that carry the genes for human telomerase reverse transcriptase (hTERT) and cyclin-dependent kinase-4 (CDK4) as described (Ramirez et al. 2004). HBEC3-KT were cultured in keratinocyte serum-free medium (Thermo Fisher Scientific, Cat # 17005042) supplemented with bovine pituitary extract as per manufacturer’s instructions (Thermo Fisher Scientific, Cat # 13028014), human growth hormone (Thermo Fisher Scientific, Cat # 1045013) and 50 U/mL penicillin/streptomycin (Thermo Fisher Scientific, Cat # 15070063).

Human foreskin fibroblasts (HFF), immortalized with hTERT (Benanti and Galloway 2004), were cultured in DMEM (Thermo Fisher Scientific, Cat # 10569010) supplemented with 10% fetal bovine serum (Thermo Fisher Scientific, Cat # 10082147) and with 50 U/mL penicillin/streptomycin (Thermo Fisher Scientific, Cat # 15070063).

All cell lines were maintained at 37°C with 5% CO_2_ and were split twice a week at a ratio of 1:4. Cells were plated in 384-well imaging plates (PhenoPlate 384-well, Revvity, Cat. # 6057500) and allowed to grow overnight until reaching approximately 80% confluency for the experiments. The seeding density per well for each cell line was optimized based on established protocols (Finn and Misteli 2021). Cells were then fixed in 4% PFA (Paraformaldehyde, Electron Microscopy Sciences, Cat # 15710) in PBS (phosphate-buffered saline, Millipore Sigma, Cat # D8537) for 10 minutes. Post-fixation, the plates underwent three PBS washes and were subsequently stored in PBS at 4°C for subsequent DNA FISH procedures.

### FISH Probes

#### DNA FISH Probes

Following established FISH protocols (Shachar et al. 2015; Hart et al. 2015; Finn and Misteli 2021), we employed BAC FISH probes RP11-717D13 and RP11-98C17 [BACPAC Resources Center] to target the downstream regions of the *MYC* or *EGFR* genes on human chromosomes 8 and 7, respectively (Supplementary Fig. 1). The generation of fluorescently labeled BAC probes involved nick translation optimized based on previous described protocols (Finn and Misteli 2021). Briefly, nick translation was performed at 14°C for 80 minutes, utilizing a reaction mixture consisting of 40 ng/mL DNA, 0.05 M Tris-HCl pH 8.0 (Thermo Fisher Scientific, Cat # 15568025), 5 mM MgCl_2_ (Quality Biological, Cat # 351-033-721), 0.05 mg/mL BSA (Millipore Sigma, Cat # A9418), 0.05 mM dNTPs (Thermo Fisher, dATP: Cat # 10216018, dGTP: Cat #1 0218014, dCTP: Cat # 10217016) including fluorescently tagged dUTP (Dyomics, DY488-dUTP: Cat. # 488-34), 1 mM β-mercaptoethanol (Bio-Rad, Cat # 1610710), 0.5 U/mL E. coli DNA Polymerase I (Thermo Fisher Scientific, Cat # EP0042), and 0.5 mg/mL DNase I (Roche, Cat # 11284932001). The reaction was halted by adding 1 μL of 0.5M EDTA (Thermo Fisher Scientific, Cat # 15575020) per 50 μL reaction volume, followed by 10 minutes heat shock at 72°C. The resulting nick-translated probe was run on a 2% agarose gel for quality control to verify successful nick translation, indicated by a smear smaller than 1 kb (Finn and Misteli 2021).

The nick-translated probe was then ethanol-precipitated to concentrate it and to remove any residual DNase and DNA polymerase activities. The probe was resuspended in a solution containing 38 ng/μL human Cot-1 DNA (Millipore Sigma, Cat # 11581074001), 256 ng/μL yeast tRNA (Thermo Fisher Scientific, Cat # AM7119), and 0.1M Sodium Acetate (Thermo Fisher Scientific, Cat # R1181) in pre-chilled (-20°C) 70% ethanol. The mixture was vortexed, spun for 1 minute at 20,000 g, and then chilled for at least 60 minutes at -20°C. Before hybridization, the mixture was spun again at 4°C for 30 minutes at 20,000 g, the supernatant was discarded, and the pellet was air-dried for 10 minutes. Finally, the probe was resuspended in the hybridization buffer, described subsequently.

As an alternative approach, we used commercially available fluorescently labelled BAC probes (Empire Genomics). For our experiment, we used the same RP11 BAC probes tagged with Green 5-Fluorescein Conjugated dUTP (Empire Genomics).

#### RNA FISH Probes

For nascent mRNA FISH targeting intron 1 of either *MYC* or *EGFR*, we used Stellaris® RNA Probes (LGC Biosearch Technologies). These probes consist of 48 single oligonucleotides, each 20 nucleotides in length labeled with Atto647N (Supplementary Fig. 1).

### Simultaneous DNA/RNA HiFISH in 384-well plates

After PFA fixation and optional storage at 4°C in PBS, cells were washed twice with PBS and permeabilized at RT for 20 minutes using 0.5% w/v saponin (Sigma Aldrich, Cat # 47036), 0.5% v/v Triton X-100 (Sigma Aldrich, Cat # X100), and 1X RNAsecure™ (Thermo Fisher Scientific, Cat # AM7006) in PBS. Following two additional PBS washes, cells were deproteinated for 15 minutes in 0.1 N HCl and neutralized for 5 minutes in 2X Saline Sodium Citrate buffer (2X SSC) (Sigma Aldrich, Cat # S6639) at room temperature. Cells were then equilibrated overnight in 50% formamide/2X SSC at 4°C.

For hybridization, we combined 4 μL of 0.4 μg of precipitated DNA probe (or commercial Empire Genomics probe) with 0.5 μL of a 12.5 μM stock Stellaris® RNA probes and 5.5 μL of hybridization buffer which is made up of 30% formamide (pH 7.0), 10% Dextran Sulfate, 0.5% Tween-20, 2X SSC, 0.5X RNAsecure™ RNAse inhibitor, and 3% THE RNA Storage Solution (Thermo Fisher Scientific, Cat # AM7001) dissolved entirely in molecular H_2_O. The hybridization mixture was shaken at 37°C for 5 minutes. Samples were washed with pre-warmed Wash Buffer (10% formamide in 2X SSC), followed by incubation at 37°C for 10 minutes.

Subsequently, 10 μL of the probe mixture was added to each well. The plate was centrifuged to eliminate bubbles, sealed, and denatured at 85°C for 7 minutes using a ThermoMixer® C-PCR 384 (Eppendorf). After denaturation, the plate was immediately transferred to a 37°C incubator for a 48-hour hybridization.

Post-hybridization, the plate was washed several times, first in Wash Buffer (10% formamide in 2X SSC) at 37°C for 1 hour, then once at RT with 2X SSC, and then with 45°C pre-warmed 1X SSC and 0.1X SSC each washed thrice for 5 minutes. Finally, DNA was stained with 3 mg/mL DAPI (4′,6-diamidino-2-phenylindole) for 10 minutes, rinsed three times with PBS, and stored in PBS until the imaging step of the protocol.

### Sequential DNA/RNA HiFISH in 384-well plates

Post PFA fixation, cells were permeabilized overnight at 4°C with pre-chilled (-20°C) 70% ethanol. Following ethanol removal, cells were washed once with wash buffer containing 10% formamide in 2X SSC at 37°C for 10 minutes. RNA hybridization was performed by adding 10 μL/well of 0.63 μM final concentration of Stellaris® RNA Probes in RNA FISH hybridization buffer (10% formamide, pH 7.0, 10% Dextran Sulfate, 2X SSC). After adding the probe mixture, the plate was centrifuged, sealed, and incubated overnight at 37°C.

Post-RNA hybridization, the plate underwent a series of washes: Wash Buffer (10% formamide in 2X SSC) at 37°C for 1 hour, followed by two consecutive room temperature washes with 2X SSC. Cells were stained with 3 mg/mL DAPI for 15 minutes, rinsed three times, and mounted in PBS for subsequent imaging on a high-throughput confocal microscope.

Post-image acquisition, a second permeabilization step was conducted using Triton/Saponin, as detailed in the Simultaneous protocol. Following overnight formamide equilibration, 4 μL of 0.4 μg of precipitated DNA probe (or specified commercial source) was resuspended in 6 μL of DNA FISH hybridization buffer (50% formamide pH 7.0, 10% Dextran Sulfate, 1% Tween-20, 2X SSC in molecular H2O). 10 μL/well of the probe mixture was added to the plate and then centrifuged and sealed. Denaturation at 85°C for 7 minutes was performed, followed by immediate transfer to a 37°C incubator for a 48-hour hybridization.

Post-hybridization, plates were rinsed once with 2X SSC at room temperature, followed by three rinses with 1X SSC and 0.1X SSC, all pre-warmed to 45°C. Cells were stained with 3 mg/mL DAPI for 10 minutes, rinsed thrice, and mounted in PBS for subsequent imaging on a high-throughput confocal microscope.

### High-throughput image acquisition

High-throughput imaging was conducted using a Yokogawa CV8000 high-throughput spinning disk confocal microscope equipped with 405 nm (DAPI Channel), 561 nm (DNA probe channel), or 640 nm (RNA probe channel) excitation lasers. A 405/488/561/640 nm excitation dichroic mirror, a 60X water objective (NA 1.2), and 445/45 nm (DAPI Channel), 525/50 nm (DNA probe channel), or 676/29 nm (RNA probe channel) bandpass emission mirrors were employed in front of a 16-bit sCMOS camera (2048 × 2048 pixels, binning 1X1, pixel size: 0.108 microns). Z-stacks spanning 7 microns were acquired at 1-micron intervals and then maximally projected in real-time.

For imaging of simultaneous DNA/RNA HiFISH, the acquisition of all three channels for DNA, RNA, and DAPI immediately followed the completion of the FISH procedure. Conversely, sequential DNA/RNA HiFISH required two separate acquisitions: the initial acquisition occurred post RNA FISH to detect the RNA FISH and DAPI signals, followed by a subsequent acquisition post DNA FISH to detect the DNA and DAPI signals (Fig. 1). Typically, between 500-5000 alleles were imaged per sample.

### Image pre-processing

For the detection of DNA and RNA signals by simultaneous hybridization, images were directly used for image analysis, as described below.

For the detection of DNA and RNA signals by sequential hybridization, the separate images generated by DNA imaging and RNA imaging required registration to align the DNA and RNA signals using the DAPI patterns. Images were subjected to an image registration algorithm based on the computation of the translation vector using cross-correlation techniques to align DNA and RNA images. The registration algorithm utilizes cross-correlation, a standard technique in signal processing, to determine the spatial translation required to align two images. In our case, these are the RNA and DNA images.

For two grayscale images *A* (representing the DNA image) and *B* (representing the RNA image), the cross-correlation *C* at a displacement (*Δx*, *Δy*) is calculated as:

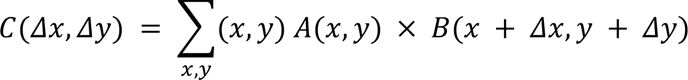

Here, *x* and *y* are the pixel coordinates in the images, and the sum is taken over all pixels where *A* and the shifted *B* overlap.

The peak of the cross-correlation function *C* indicates the displacement at which the images are best aligned. The coordinates of this peak (*Δx*_*peak*_, *Δy*_*peak*_) represent the translation vector.

Once the translation vector (*Δx*_*peak*_, *Δy*_*peak*_) is determined, it is applied to the RNA image to achieve alignment with the DNA image.

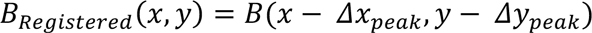

In this equation, *B*_*Registered*_ is the spatially shifted RNA image, and the operation ensures that every pixel in *B* is moved according to the calculated translation vector. Boundary pixels are set to zero.

The image registration code is publicly available on GitHub: https://github.com/CBIIT/DNA_RNA_registration

### High-throughput Image Analysis

After image acquisition of simultaneous DNA/RNA FISH or after registration of separate RNA and DNA images for sequential FISH, we employed High-Throughput Image Processing Software (HiTIPS) (Keikhosravi et al. 2023) to analyze the image dataset consisting of DNA, RNA, and DAPI channels. Nuclei were segmented using the HiTIPS’s GPU-accelerated implementation of the CellPose deep learning segmentation model (Stringer et al. 2021), along with the Laplacian of Gaussian method for detecting FISH signals as described in (Keikhosravi et al. 2023). Before starting the analysis, these parameters were adjusted using real-time visual feedback provided by overlaying the results of the segmentation on the original images to maximize segmentation accuracy. Each plate was analyzed using specifically chosen parameters such as average cell size, Laplacian of Gaussian kernel size, thresholding method, etc., to maximize spot detection accuracy.

To calculate the radial position of FISH signals within the nucleus, first, a distance transform was calculated from binary images of nuclei. The distance transform returns the closest distance of each pixel from the background, meaning the pixels at the boundary have a value of 0, and the pixels at the center of the nucleus have the maximum value. The distance transform of each nucleus is then normalized separately and subtracted from 1. This will return 1 for the pixels at the periphery of the nucleus, which is farthest from the nucleus center, and 0 for the pixels at the center.

Single-cell and single-spot results were separately saved as flat text files for downstream data analysis. All image processing was done on the NIH HPC BioWulf cluster (https://hpc.nih.gov/) to maximize spot detection accuracy.

### Data analysis

Data analysis was performed using R (R Core Team 2023) and these R packages: tidiverse (Wickham et al. 2019), SpatialTools (Joshua French 2023), fs (Hester et al. 2023), reshape2 (Wickham 2007), data.table (Barrett et al. 2024), and ggthemes (Arnold et al. 2024).

Briefly, single-cell data were read from several flat text files (One file per well) output by HiTIPS and concatenated in a single data frame for each experiment. Cells with nuclei smaller than 10 microns and a circularity value < 0.95 were filtered out as segmentation errors. FISH spot-level data were read from several flat text files (One file per well and per channel) output by HiTIPS and concatenated in a single data frame per experiment. Spot level data contain information on which nucleus/cell each spot belongs to. Spots that did not overlap with any nucleus in the image were filtered out.

Based on the spot level data, we calculated the number of spots for each cell and each channel. These discrete numerical values were further binned in a new variable for each channel that assumed the “0”, “1”, “2” and “>=3" values. Only cells with 2 DNA FISH spots and 2 or less RNA FISH spots were analyzed for downstream DNA FISH/RNA FISH Euclidean distances, and for DNA FISH radial distance calculations or plots.

We used the X and Y coordinates for each spot to first calculate all the possible distances between the 2 DNA FISH spots and RNA FISH spots, if any, on a per-cell basis. We then calculated the minimum DNA FISH/RNA FISH Euclidean distance on a per DNA FISH spot basis. DNA FISH spots in cells that did not contain an RNA FISH spot were automatically assigned an “NA” value for the minimum DNA FISH/RNA FISH Euclidean distance. DNA FISH spots with a value of “NA” were classified as “No Transcription.” DNA FISH spots with a DNA FISH/RNA FISH Euclidean distance below 1 micron, indicating proximity to an RNA FISH spot, were classified as “Active.” Otherwise, the remaining DNA FISH spots were classified as “Inactive.” In addition, normalized radial distances, which can assume a continuous value between 0 and 1, were binned into “shells” at intervals of 0.2.

Normalized radial distance distributions between the active vs. inactive in simultaneous and the sequential protocols were compared using a two-sided Kolmogorov-Smirnov test.

Plots and tables from single experiments were generated by R, compiled in Microsoft Excel, and then organized into figures using BioRender.com.

No primary human or animal samples were used in these studies.

## Results

### High-throughput detection of DNA and RNA

We established two separate high-throughput FISH pipelines for the combined detection of DNA and RNA at individual alleles. In the first approach, DNA and RNA are visualized simultaneously using a single hybridization reaction using a mixture of DNA and RNA probes. The DNA and RNA signals are then imaged at the same time (Fig. 1). In a second approach, RNA and DNA are detected sequentially in two separate hybridization reactions, and imaging of RNA and DNA, respectively, occurs after each hybridization step (Fig. 1). In both approaches, we utilized chromosome-specific BAC FISH probes synthesized following established protocol (Finn and Misteli 2021) in conjunction with commercially designed Stellaris® RNA FISH probes (Orjalo et al. 2011). These methodologies share similarities in cell handling and final image processing and analysis. However, they differ in the cell permeabilization, probe hybridization, and image acquisition stages (Fig. 1).

### Simultaneous DNA/RNA HiFISH

We sought to develop a protocol for the sensitive visualization of cellular DNA and RNA through single-step hybridization of a mixture of DNA and RNA probes (Fig. 1). To do so, we optimized all steps of the FISH protocol, including permeabilization, hybridization, and washes in a 384-well plate format (Fig. 1; see Materials and Methods).

For optimal results, cells were grown to 80% density, and standard 4% paraformaldehyde fixation was used. For cell permeabilization, a saponin/triton combination supplemented with an RNase inhibitor was used to preserve RNA integrity. After deproteination with HCl, the cells were equilibrated overnight in 50% formamide in preparation for subsequent hybridization.

The hybridization step employed a customized buffer designed for optimal DNA and RNA probe binding, containing dextran sulfate, formamide, SSC, Tween-20, sodium citrate, and an RNase inhibitor (see Materials and Methods for details). The concentration of the components in the hybridization buffer was systematically adjusted in pilot experiments to an intermediate level between standard DNA and RNA hybridization buffers (Shaffer et al. 2013; Finn and Misteli 2021), creating an environment conducive to both DNA and RNA probe hybridization. The hybridization buffer was explicitly formulated for simultaneous DNA/RNA HiFISH, with the formamide concentration adjusted to be 20% lower than that typically used in DNA FISH protocols, yet 20% higher than that in RNA FISH, to balance the conditions for both types of probes. Moreover, we halved the concentration of Tween-20 to approximate the viscosity characteristic of RNA FISH buffers. Dextran sulfate and SSC concentrations were kept at standard levels compatible with both DNA and RNA FISH. This tailored approach ensures the stability and hybridization efficacy of both DNA and RNA probes. Additionally, the buffer was supplemented with an RNase inhibitor, and a molecular grade 1 mM sodium citrate buffer was used, aimed at stabilizing and minimizing inherent DNA and RNA base hydrolysis by inhibiting the activity of both DNase and RNase during the hybridization process. Detection was further optimized using a pre-hybridization wash buffer containing 10% formamide and 2XSSC. We either used commercially available pre-labeled DNA probes or in-house generated fluorescently labeled BAC probes (see Materials and Methods). For RNA probes, we used multiple 20-base single-stranded DNA oligonucleotides, individually labeled and designed to bind distinct segments of the target RNA via Watson-Crick base pairing (Supplementary Fig. 1).

During the process of optimizing the simultaneous FISH protocol to a 384-well format, we noted heterogeneity in heat distribution among wells during the heat denaturation step, especially when using various commercial heat blocks. Because uneven heating across the plate could introduce inconsistencies in technical replicates, we systematically assessed multiple heat blocks. We identified the ThermoMixer® C-PCR 384 (Eppendorf) as the optimal choice, providing a reliable and uniform heat distribution across the 384-well plate.

Hybridization efficiency was further optimized by a prolonged incubation period of up to 48 hours. All steps used molecular-grade RNase-free reagents, and treatment of reagents with RNAsecure™ ensured RNase inactivation and thorough RNase decontamination measures were implemented on equipment and benchtops using RNaseZap™ before experiments. Commercially available or in-house labeled BAC probes worked equally well, and Stellaris® RNA FISH probes were routinely used. Following a sequential series of standard rinsing steps, cells are stained with DAPI and mounted for imaging using a high-throughput confocal microscope per standard FISH protocol (see Materials and Methods).

To evaluate our approach, we performed simultaneous DNA/RNA HiFISH utilizing BAC FISH probes targeting the downstream regions of the *MYC* and *EGFR* genes on human chromosomes 8 and 7, respectively (Supplementary Fig. 1). We detected both DNA and RNA for *MYC* and *EGFR* in HBEC and HFF cells with high efficiency (Fig. 2, 3; Supplementary Fig. 2, 3). *MYC* DNA signals were detected in 98 ± 1.1% of HBECs and 99 ± 0.9% of HFF cells, and both copies of *MYC* were detected in 72 ± 5.7% of HBEC cells (Fig. 2c) and 77 ± 2.9% of HFF cells (Fig. 2i). Likewise, *EGFR* was detected in 89 ± 4.4% of HBECs and 90 ± 9.2% of HFFs, 57 ± 7.6% of HBEC cells (Fig. 3c), and 58 ± 18% of HFF cells showing two signals (Fig. 3i). These values are well within detection efficiencies previously reported in hiFISH approaches (Shachar et al. 2015; Finn et al. 2019). The few DNA signals that were missed are likely due to suboptimal FISH hybridization, weak FISH signal, or high background signal, which can reduce signal detection.

**Fig. 2.**
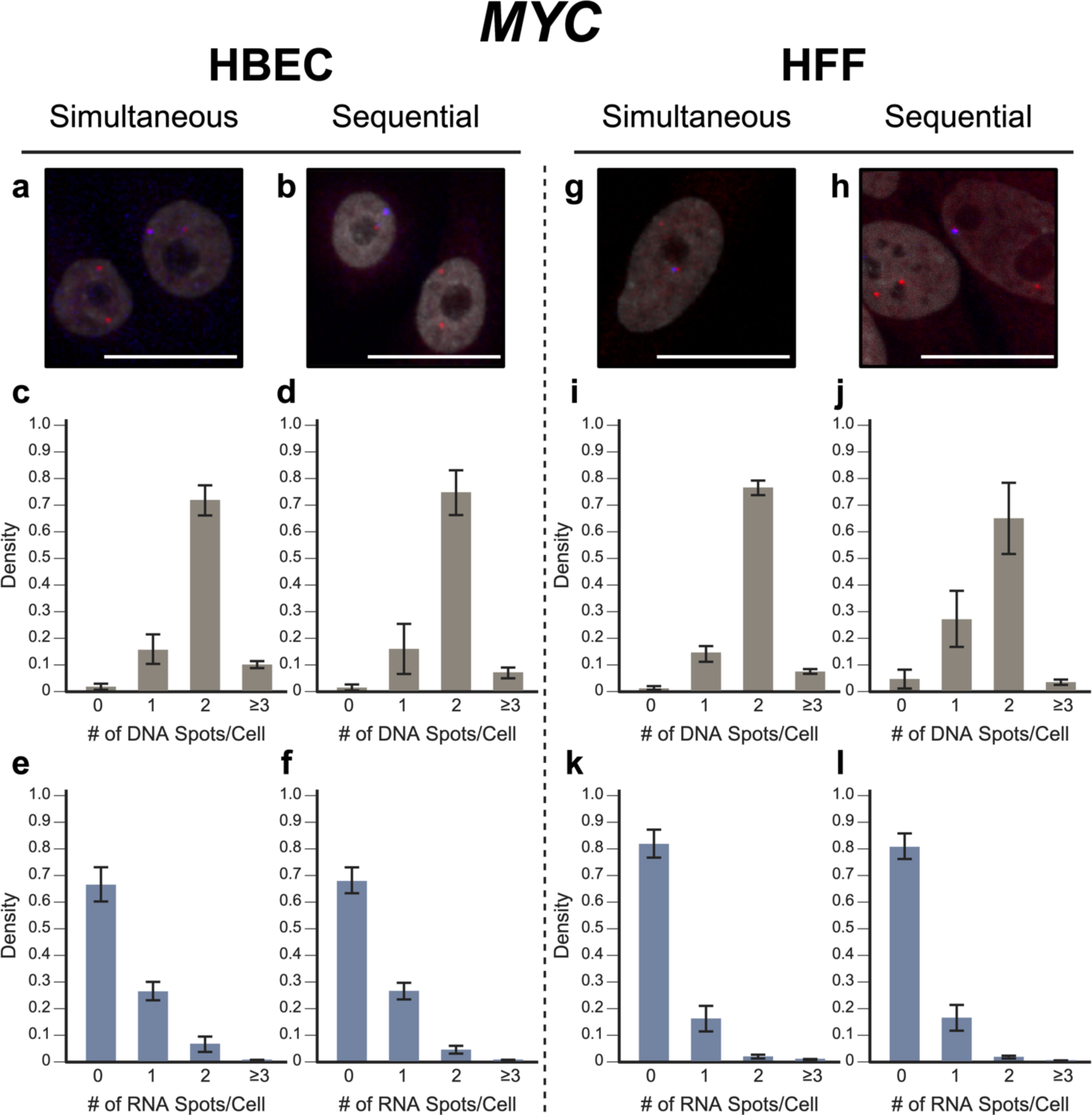
MYC simultaneous vs. sequential DNA/RNA HiFISH detection. (a-b) Representative images of simultaneous and sequential MYC DNA/RNA HiFISH in HBEC cells (red: DNA, blue: nascent mRNA, grey: DAPI-stained nucleus). (c-d) Histograms of the number of detected DNA signals per nucleus in HBEC cells. 10,432 alleles for simultaneous and 7,726 alleles for sequential detection were analyzed. (e-f) Histograms of nascent mRNA signals per nucleus in HBEC cells. 10,432 alleles for simultaneous and 7,726 alleles for sequential detection were analyzed. (g-h) Representative images of simultaneous and sequential MYC DNA/RNA HiFISH in HFF cells. (i-j) Histograms of the number of DNA signals per nucleus in HFF cells. 4,657 alleles for simultaneous and 4,126 alleles for sequential detection were analyzed. (k-l) Histograms of nascent mRNA signals per nucleus in HFF cells. 4,657 alleles for simultaneous and 4,126 alleles for sequential detection were analyzed. All values represent means ± s.d. of at least two independent experiments. Scale bars: 20 µm

**Fig. 3.**
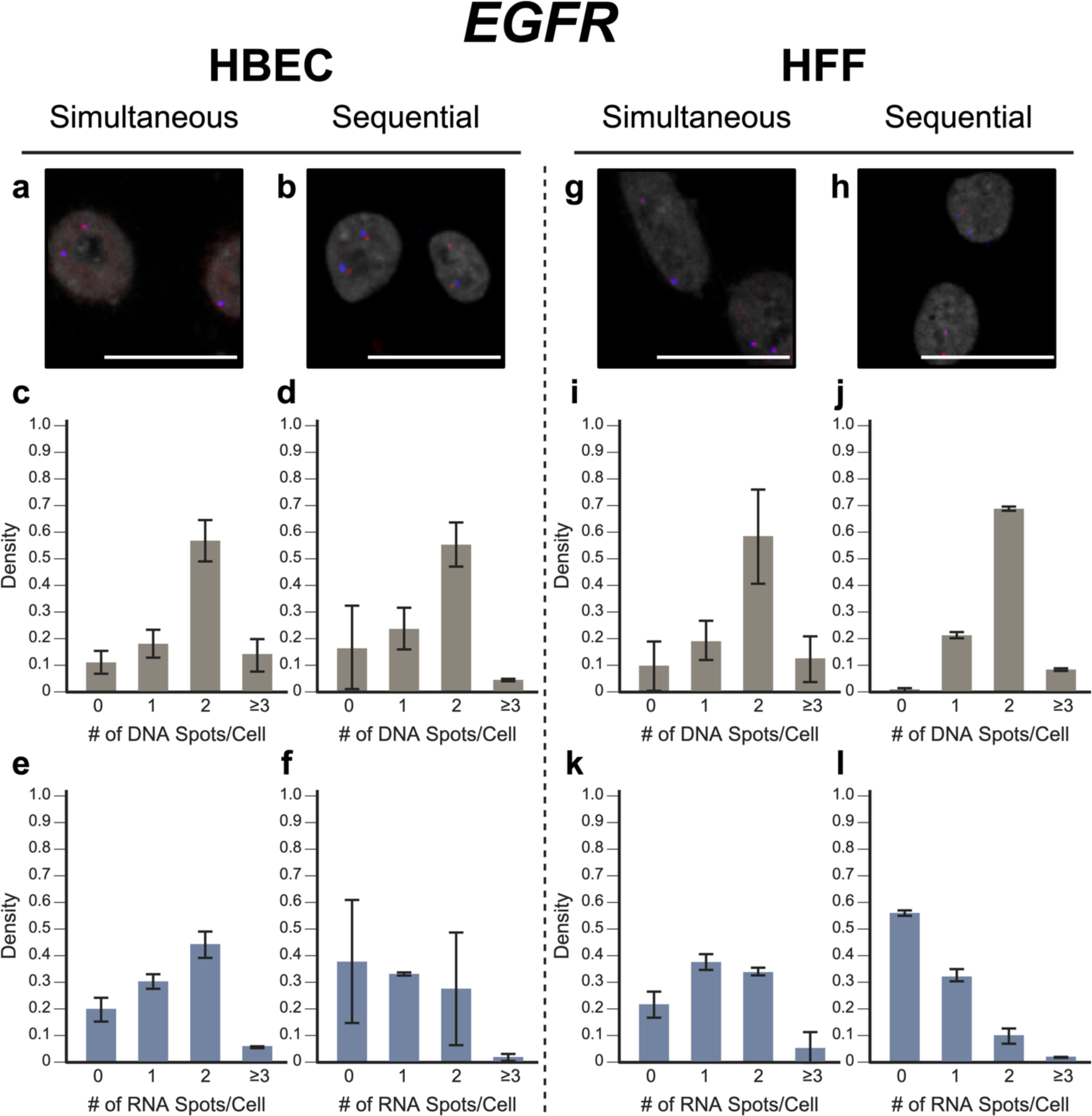
EGFR simultaneous vs. sequential DNA/RNA HiFISH detection. (a-b) Representative images of simultaneous and sequential EGFR DNA/RNA HiFISH in HBEC cells (red: DNA, blue: nascent mRNA, grey: DAPI-stained nucleus). (c-d) Histograms of the number of detected DNA signals per nucleus in HBEC cells. 23,740 alleles for simultaneous and 5,403 alleles for sequential detection were analyzed. (e-f) Histograms of nascent mRNA signals per nucleus in HBEC cells. 23,740 alleles for simultaneous and 5,403 alleles for sequential detection were analyzed. (g-h) Representative images of simultaneous and sequential EGFR DNA/RNA HiFISH in HFF cells. (i-j) Histograms of the number of DNA signals per nucleus in HFF cells. 9,133 alleles for simultaneous and 798 alleles for sequential detection were analyzed. Data shown in (j) represents mean values ± s.d. of two technical replicates from a single experiment. (k-l) Histograms of nascent mRNA signals per nucleus in HFF cells. 9,133 alleles for simultaneous and 798 alleles for sequential detection were analyzed. Data shown in (l) represents mean values ± s.d. of two technical replicates from a single experiment. Unless indicated otherwise, values represent mean ± s.d. of at least two independent experiments. Scale bars: 20 µm

For RNA detection, in line with the demonstrated variable expression of *MYC* in individual cells (Liu et al. 2023), 6.4 ± 2.9% of HBECs showed biallelic expression, monoallelic expression in 26 ± 3.4% of cells and no expression in 66 ± 6.3% of HBEC cells (Fig. 2e). Similarly, in HFF, 2.0 ± 0.6% showed biallelic expression, monoallelic expression in 16 ± 4.8% and no expression in 82 ± 5.1% of cells (Fig. 2k). In contrast, *EGFR* RNA is more highly expressed in both HBEC and HFF and accordingly more active alleles were detected in HBEC; 44 ± 5.0% of cells showed biallelic expression, 30 ± 2.7% monoallelic expressed, and only 20 ± 4.5% no expression. In HFF, 34 ± 1.7% of cells expressed *EGFR* biallelically, 38 ± 3.1% monoallelically expressed, and 22 ± 5.0% did not express the gene. More than the expected two transcription sites per nucleus were detected in only 3.2 ± 3.1% of cells for both genes and cell lines due to false positive detection of FISH signals. We conclude that simultaneous detection of DNA and RNA using a single-hybridization step is feasible.

### Sequential DNA/RNA HiFISH

To complement the simultaneous FISH detection, we developed an alternative in which RNA and DNA are detected in a sequential fashion based on previously published protocols for each nucleic acid species (Raj and Tyagi 2010; Shaffer et al. 2013; Finn and Misteli 2021; Finn et al. 2022) with optimizations implemented for high-throughput image processing (Fig. 1). This method involves first RNA detection, imaging of the RNA signals, followed by DNA FISH and a second round of imaging to detect DNA signals. The two sets of images are then accurately superimposed using an image registration algorithm. In our hands, detection of DNA prior to RNA yielded sub-optimal results, and we focused on optimizing protocols that detect RNA before DNA.

After cell culture and fixation, the RNA FISH protocol is initiated with overnight permeabilization at 4°C using 70% ethanol. The subsequent hybridization involves washing with 10% formamide buffer, followed by the introduction of RNA probes in a standard hybridization buffer (Shaffer et al. 2013). After overnight hybridization, a series of rinsing steps is performed, and cells are stained with DAPI for imaging of RNA signals.

After imaging of the RNA signals, plates are returned for DNA FISH, which involves re-permeabilization with saponin/triton, deproteination with HCl, and equilibration in 50% formamide, followed by hybridization with DNA BAC probes in standard hybridization buffer and washes as per established protocols (Shachar et al. 2015; Finn et al. 2019) and finally imaging of the DNA signal by high-throughput microscopy.

After imaging of the DNA signals, an image alignment algorithm is employed for DAPI-stained DNA and RNA image registration, ensuring precise spatial alignment of the RNA and DNA signals for combined analysis of DNA and RNA signals (see Materials and Methods for details). The algorithm utilized a technique akin to aligning two complex patterns by identifying reference points within each image. In this case, it scrutinizes the distinct DAPI staining patterns of individual nuclei present in both the DNA and RNA images, relying on specific high-intensity areas—corresponding to the DAPI-stained regions—as reference points for alignment. By evaluating these identifiable features in RNA and DNA FISH images and leveraging cross-correlation techniques, the algorithm calculated the optimal translation vector, essentially a set of instructions determining the precise shift needed to match the RNA image to the DNA image. This alignment process enabled us to generate an accurate combined image of the RNA and DNA signals, allowing us to examine and compare the DNA and RNA signals (Fig. 2, 3; Supplementary Fig. 2, 3; see Materials and Methods for details).

In validation experiments, sequential RNA/DNA HiFISH using BAC FISH probes targeted *MYC* and *EGFR*. High detection efficiency was observed for both DNA and RNA signals in HBEC and HFF cells (Fig. 2, 3; Supplementary Fig. 2, 3). *MYC* DNA signals were detected in 98 ± 1.3% of HBEC and 95 ± 3.6% of HFF cells, with 75 ± 8.6% of HBEC cells (Fig. 2d) and 65 ± 13.5% of HFF cells (Fig. 2j) showing two copies of *MYC*. *EGFR* was detected in 83 ± 15.6% and 99 ± 0.6% of HBECs and HFFs, respectively, with 56 ± 8.1% of HBEC cells (Fig. 3d) and 69 ± 1.0% of HFF cells (Fig. 3j) showing two signals.

For *MYC* RNA detection, 27 ± 3.2% of HBECs exhibited monoallelic expression, 4.6 ± 1.5% showed biallelic expression, and 68 ± 4.9% showed no expression (Fig. 2f), similar to the simultaneous detection method. For HFF, 81 ± 4.8% of cells did not express *MYC*, 16 ± 4.8% showed monoallelic expression and 2.0 ± 0.2% showed biallelic expression (Fig. 2i). Conversely, *EGFR* RNA was more highly expressed in both HBEC and HFF. In HBEC, 27 ± 21% of cells showed biallelic expression, 33 ± 0.6% monoallelic expressed, while 38 ± 23% of cells were silent (Fig. 3f). In HFF, 10 ± 3.0% of cells were biallelic expressed, 33 ± 2.2% were monoallelic expressed, and 56 ± 1.0% of cells were silent (Fig. 3i).

When directly compared, simultaneous and sequential DNA/RNA HiFISH gave similar results but exhibited some nuanced differences. Simultaneous HiFISH was slightly more robust in detecting *MYC* DNA signals, with detection rates of 98-99% in HBECs in HFF cells vs. 95-985%, in sequential FISH. The detection efficiency of *EGFR* DNA signals was similar for both methods. In terms of RNA detection, *MYC* RNA displayed comparable expression in simultaneous and sequential detection of *MYC* active alleles (33 ± 6.2% vs 31 ± 4.8% in HBECs, 18 ± 5.3% vs 18 ± 5.0% in HFFs, respectively). In contrast, detection of the more highly expressed *EGFR* RNA by simultaneous FISH was slightly more efficient compared to sequential detection with detection efficiencies in HBEC of 74 ± 4.5% vs 61 ± 22%, and in HFFs 72 ± 1.4% vs 42 ± 0.7%. As expected, differences between the FISH methods were more pronounced for the more highly expressed *EGFR* gene, whereas the two methods were more similar in detection efficiency for the more lowly expressed *MYC* gene. Considering the detection sensitivities, we conclude that both methods are suitable for the high-sensitivity detection of DNA and RNA at the single-allele level.

### Application of allele-level DNA/RNA HiFISH to compare the radial position of active and inactive gene alleles

The RNA/DNA detection pipelines developed here can be used to probe the behavior of active and inactive alleles in the same cell nucleus. As proof-of-principle for their utility, we applied DNA/RNA HiFISH to ask whether the nuclear position of the active and inactive alleles differ (Fig. 4). To do so, we visualized active and inactive alleles of *MYC* or *EGFR* in HBEC or HFF using both simultaneous and sequential DNA/RNA HiFISH. We defined *active alleles* as DNA FISH signals associated with an RNA signal within 1.0 micron. DNA FISH signals without an RNA signal were defined as inactive. For each active or inactive allele, we determine its radial position relative to the center of the cell nucleus using a previously described distance transform method, which assigns each allele a value between 0 (center of the nucleus) and 1 (periphery) (see Materials and Methods for details). We analyzed between 1,280 and 5,118 alleles per sample. To ensure the reliability of our analyses, we excluded from analysis nuclei that contained an incorrect number of detected signals for both DNA and RNA, typically under 10–30% nuclei in the sample.

**Fig. 4.**
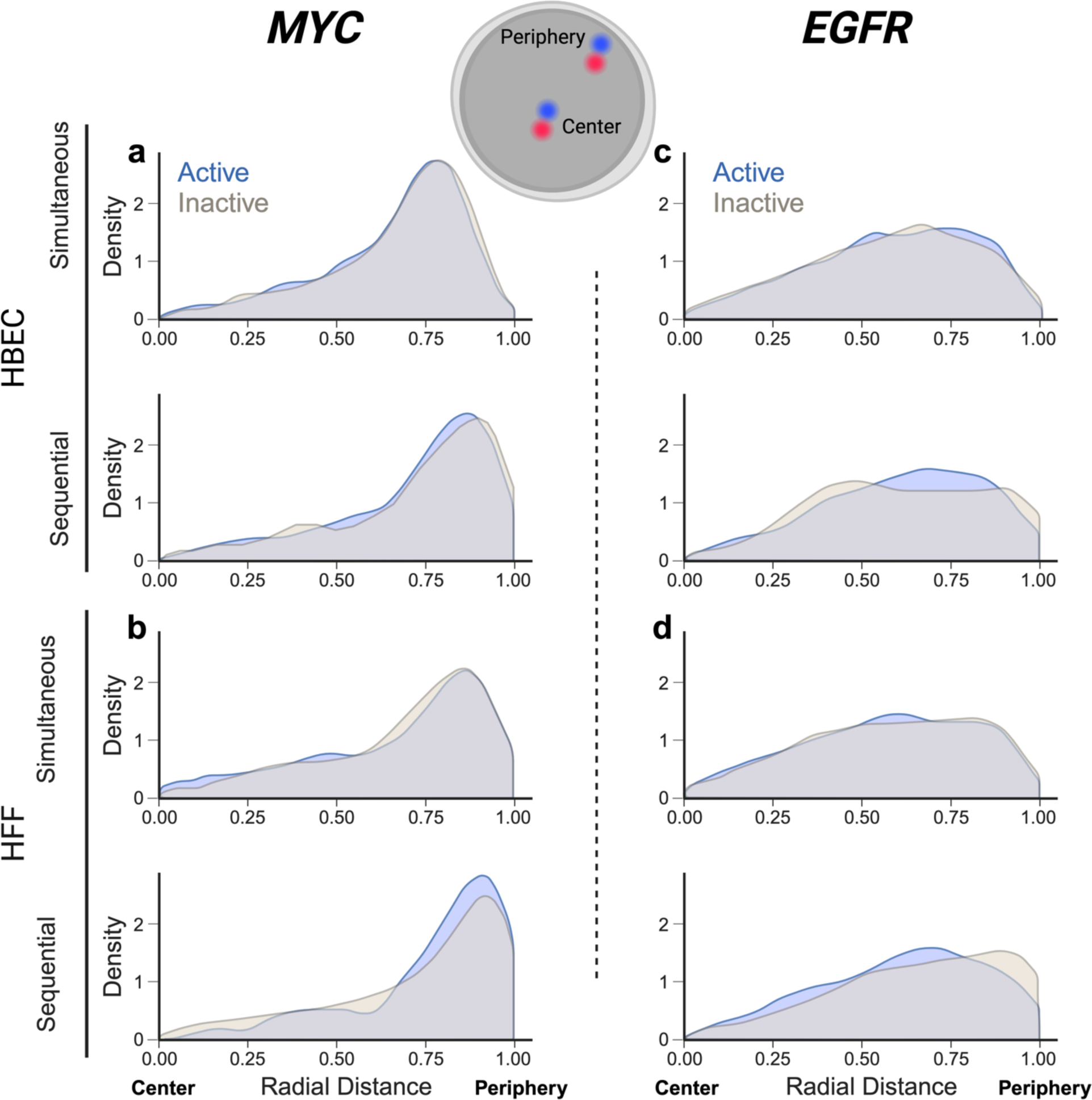
Radial position distribution of active vs inactive alleles. (a – b) Radial distribution for active or inactive MYC alleles: (a) in HBEC cells, 1,914 alleles for simultaneous and 1,380 alleles for sequential detection were analyzed. (b) in HFF cells (490 alleles for simultaneous, 790 alleles for sequential detection). (a – b) Radial distribution for active or inactive EGFR alleles: (a) in HBEC cells (4,614 alleles for simultaneous, 504 alleles for sequential detection). (b) in HFF cells (3,458 alleles for simultaneous and 368 alleles for sequential detection). Values represent a single representative dataset from 2-3 experiments, except for sequential detection of EGFR in HFF, which is a single experiment.

In line with prior observations (Shachar et al. 2015), the overall distribution of the radial position for the two genes was distinct (Fig. 4). *MYC* showed preferential localization to the periphery of the nucleus with both simultaneous and sequential methods in both HBEC (mean radial distance = 0.7 ± 0.1) and HFF (mean radial distance = 0.7 ± 0.0). This observation aligns with previous findings indicating peripheral localization of *MYC* in normal HCECs and cancer HCT-116 cells (Scholz et al. 2019). In contrast, *EGFR* exhibited a more uniform distribution within the nucleus in both HBEC (mean radial distance = 0.6 ± 0.0) and HFF (mean radial distance = 0.6 ± 0.0) (Fig. 4 e - h).

A comparison of the radial position of active vs. inactive alleles for *MYC* and *EGFR* showed no difference in localization. For *MYC* active and inactive alleles in HBEC showed similar peripheral positioning using both simultaneous (mean radial distance = 0.7 ± 0.2 for both) or sequential DNA/RNA HiFISH (KS test p-values: 0.37, 0.22, respectively) (Fig. 4; Supplementary Table 1). Similarly, the radial positioning of *MYC* in HFF showed similar peripheral positioning for both active (mean radial distance = 0.7 ± 0.3) and inactive (mean radial distance = 0.7 ± 0.2) alleles using either simultaneous FISH or sequential detection (mean radial distance = 0.7 ± 0.2) (KS test p-values: 0.86, 0.13, respectively) (Fig. 4; Supplementary Table 1).

Like *MYC*, the radial positioning of *EGFR* did not show a difference between active and inactive alleles in both HBEC and HFF (mean radial distance = 0.6 ± 0.2; KS test p-value: 0.51± 0.2) with the exception of inactive alleles in HFF when using the sequential method (mean radial distance = 0.7 ± 0.2) which may be attributed to smaller sample size (KS test p-value: 0.05) (Fig. 4; Supplementary Table 1).

Comparing the radial distance distributions obtained by simultaneous versus sequential DNA/RNA HiFISH, we observed a slight but statistically significant difference, with an average KS test p-value of 0.04 ± 0.06 (Supplementary Table 2). Sequential FISH exhibited a mean radial distance skew of 0.05 ± 0.02 units towards the nuclear periphery compared to simultaneous FISH (Fig. 4; Supplementary Table 2). We suspect this discrepancy may arise from cell shrinkage during sequential FISH due to the use of a 70% ethanol overnight permeabilization step.

Together, these results demonstrate sensitive detection of active and inactive gene alleles and suggest that the radial position of *MYC* and *EGFR* alleles is independent of their expression status.

## Discussion

In this study, we have developed and optimized two high-throughput DNA/RNA FISH protocols, one for simultaneous detection and the other for sequential detection of DNA and RNA in individual cell nuclei. These protocols, implemented in a 384-well plate format alongside automated image analysis, provide a versatile and sensitive tool for mapping chromatin features and gene activity at the single-allele level and at high throughput. To our knowledge, this is the first description of a simultaneous DNA/RNA FISH detection protocol, regardless of the degree of throughput.

To optimize our detection methods, we systematically fine-tuned each protocol step, including cell permeabilization, hybridization, and choice of hybridization buffers. In simultaneous FISH, as previously reported (Lai et al. 2013), we encountered challenges due to the fragile nature of RNA molecules. Intrinsically, mRNA targets are prone to damage under the stringent conditions needed for DNA FISH, such as high temperatures and low pH, which are necessary to denature the target DNA for detection. Consequently, it was imperative to ensure an RNase-free and RNA-protective microenvironment by using RNase inhibitors and buffers with low pH and efficient chelating properties. It was also essential to maintain hybridization conditions that effectively supported both DNA and RNA hybridization by adjusting the key components of the hybridization buffer to concentrations between those typically used in DNA and RNA hybridization protocols. In addition, we used a high concentration of RNA oligonucleotide probes, and we took measures to ensure accurate heat distribution during denaturation. For sequential FISH, incorporating a permeabilization step in each detection protocol was critical for the success of the DNA FISH. These optimizations were crucial for achieving high detection efficiency for both DNA and RNA signals.

One of the key advantages of our methods is the ability to perform high-throughput imaging, enabling the analysis of a large number of cells and individual alleles. We routinely image several thousand cells per sample, yielding high statistical power in these single-cell analyses. These approaches are complementary to traditional biochemical approaches, which often provide population averages and lack single-cell resolution (Akgol Oksuz et al. 2021). Using high-throughput FISH approaches is especially valuable when studying single-cell heterogeneity, as it allows the capture of a representative sample of cells and drawing meaningful conclusions. The use of 384-well plates and automated image analysis streamlines the process and ensures reproducibility. We demonstrate high efficacy and sensitivity of both methods by successfully visualizing *MYC* and *EGFR* DNA and RNA in multiple cell types. We routinely detect more than 95% of expected DNA signals. We also demonstrate reliable RNA detection. In line with the recent realization that all human genes undergo stochastic cycles of gene activity and inactivity, we find cells with zero, one, or two active alleles in the same nucleus. Our DNA/RNA visualization approach indicates that at any given time, more *EGFR* alleles are active than *MYC* alleles. This difference corresponds to the higher expression level of *EGFR* than *MYC* in the cell lines analyzed here.

We applied the newly developed DNA/RNA HiFISH methods to probe the radial position of genes. In line with previous studies that have reported distinct radial positioning patterns for individual genes (Shachar et al. 2015; Scholz et al. 2019), we observed distinct radial positioning patterns for *MYC* and *EGFR*, with *MYC* alleles showing a preference for peripheral localization, while *EGFR* alleles exhibited a more uniform distribution with a slight preference for the periphery. Our DNA/RNA HiFISH methods are particularly well suited to address the long-standing question of whether active genes occupy a distinct nuclear location from inactive genes. While some examples exist of a correlation between gene activity and nuclear location, such as the internalization of IgH and Igkappa during lymphocyte development (Kosak et al. 2002), the majority of studies do not support any correlation between gene activity and location (Meaburn and Misteli 2008; Nakayama et al. 2022). However, these studies relied entirely on DNA FISH, and gene activity status was assessed on a population-wide basis rather than at the single-allele level. Our combined DNA/RNA HiFISH allowed us to extend these studies and probe the relationship between gene activity and location at the allele level in individual nuclei. Using this highly sensitive approach, we do not find differential locations of active vs inactive alleles for *MYC* and *EGFR*. This observation agrees with a recent analysis of the location of active *MYC* alleles, which found them to be located both at the nuclear periphery and the interior (Scholz et al. 2019). Interestingly, these findings on two bi-allelically expressed genes differ from that reported for the mono-allelically expressed *GFAP* gene in astrocytes, where the inactive allele is more frequently found towards the periphery and the active allele more internally (Takizawa et al. 2008). Although the sample size is small, these findings may suggest that stochastically silent gene alleles do not change their nuclear location; epigenetically altered alleles, however, may.

In conclusion, our optimized DNA/RNA HiFISH protocols provide new tools for studying the relationship between chromatin structure and gene activity at the single-allele level and at high throughput. The ability to capture single-cell heterogeneity and perform large-scale analyses makes this a valuable method for investigating the role of nuclear positioning and chromatin organization in gene regulation.

## Acknowledgments

We thank members of the Misteli laboratory and the NCI High-Throughput Imaging Facility for discussions and input. Computation was performed on the NIH HPC Biowulf cluster. F.A. was supported by a graduate fellowship from the Ministry of Education of Saudi Arabia. Work in the Misteli Lab is supported by the Intramural Research Program of the NIH, NCI, Center for Cancer Research through grant 1-ZIA-BC010309-24 and as part of the 4D Nucleome Common Fund.

## Supplementary Information

**Supplementary Figure 1:**
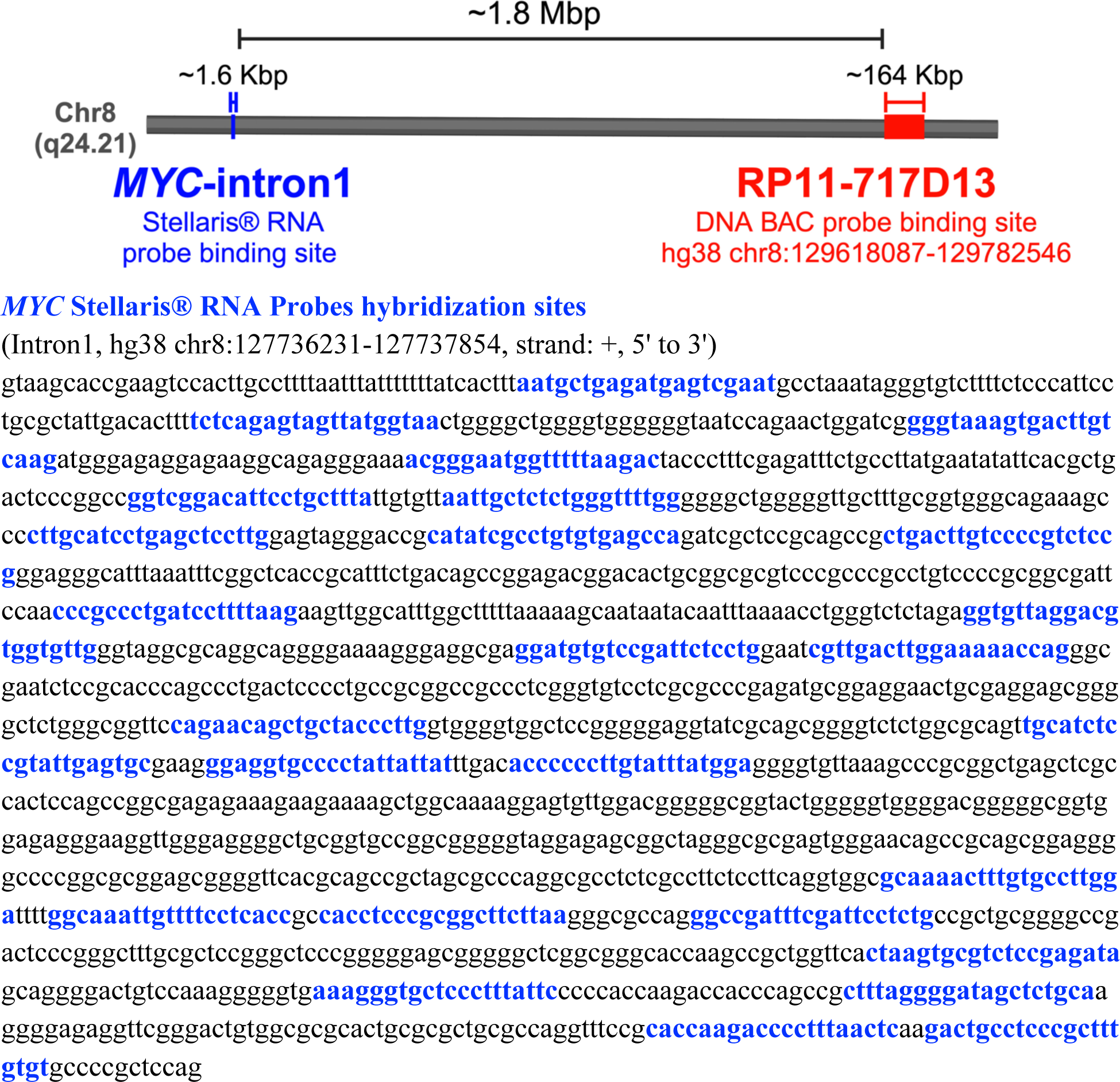
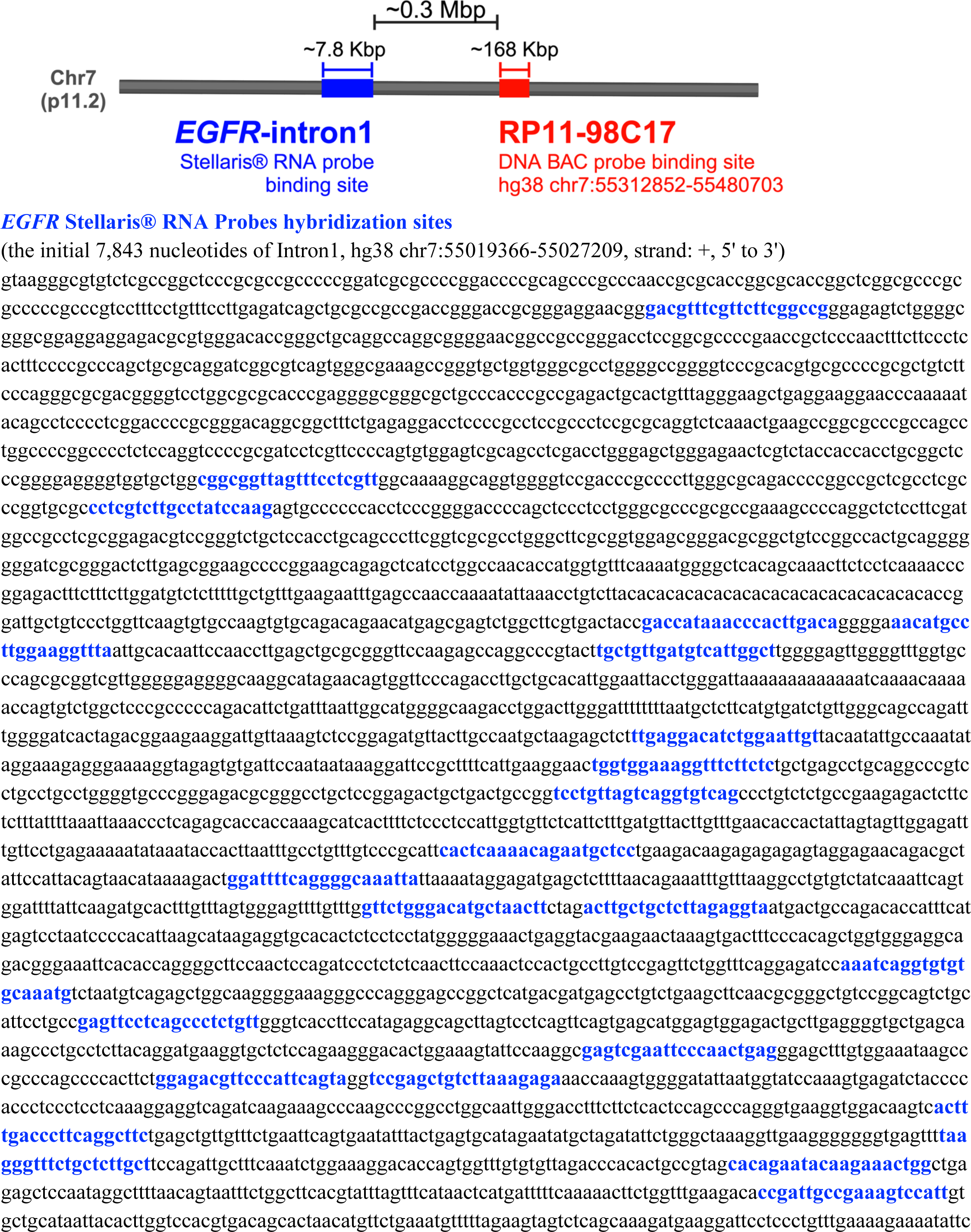
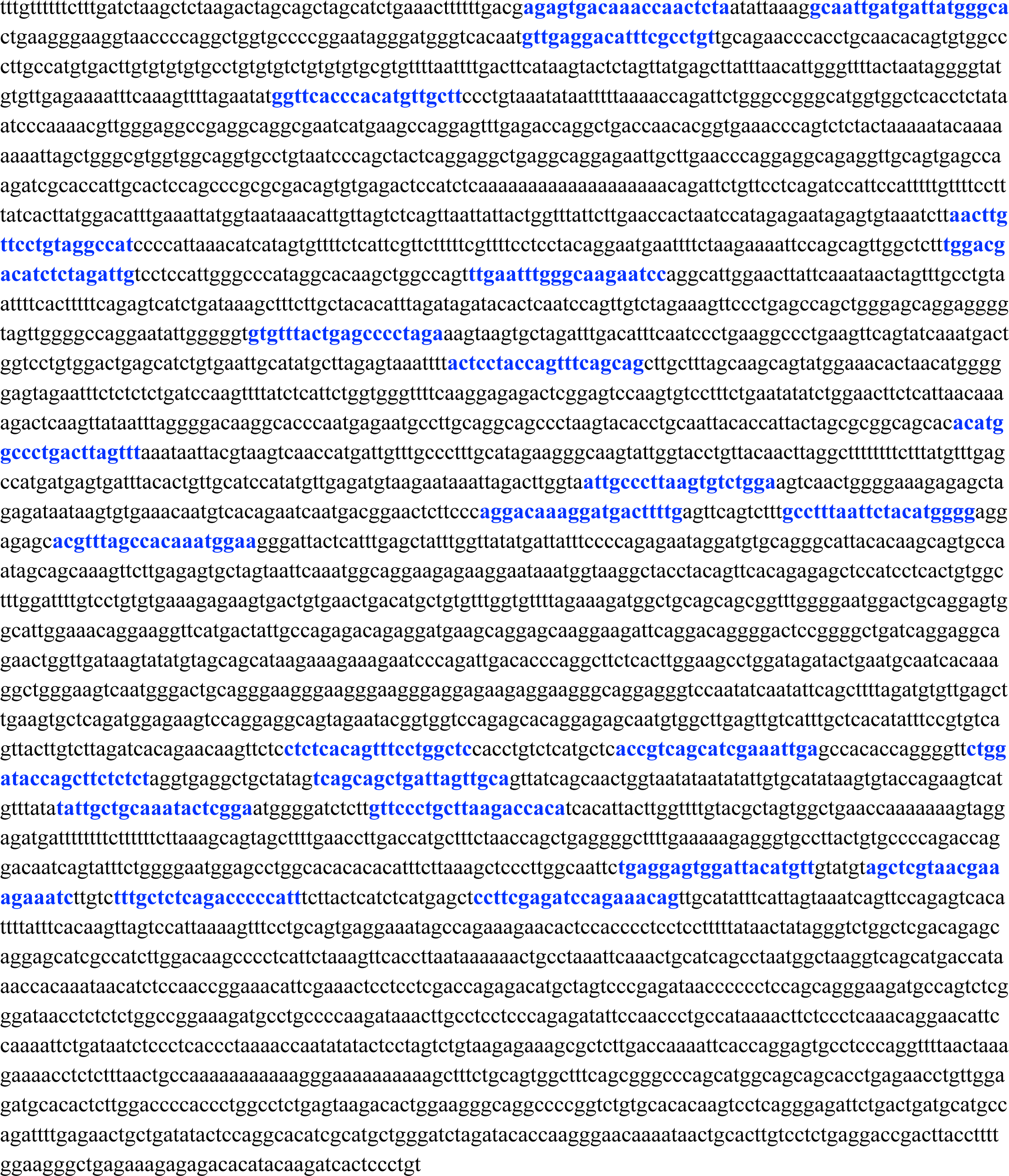
Sequence and location of DNA and RNA probes binding sites.

**Supplementary Figure 2:**
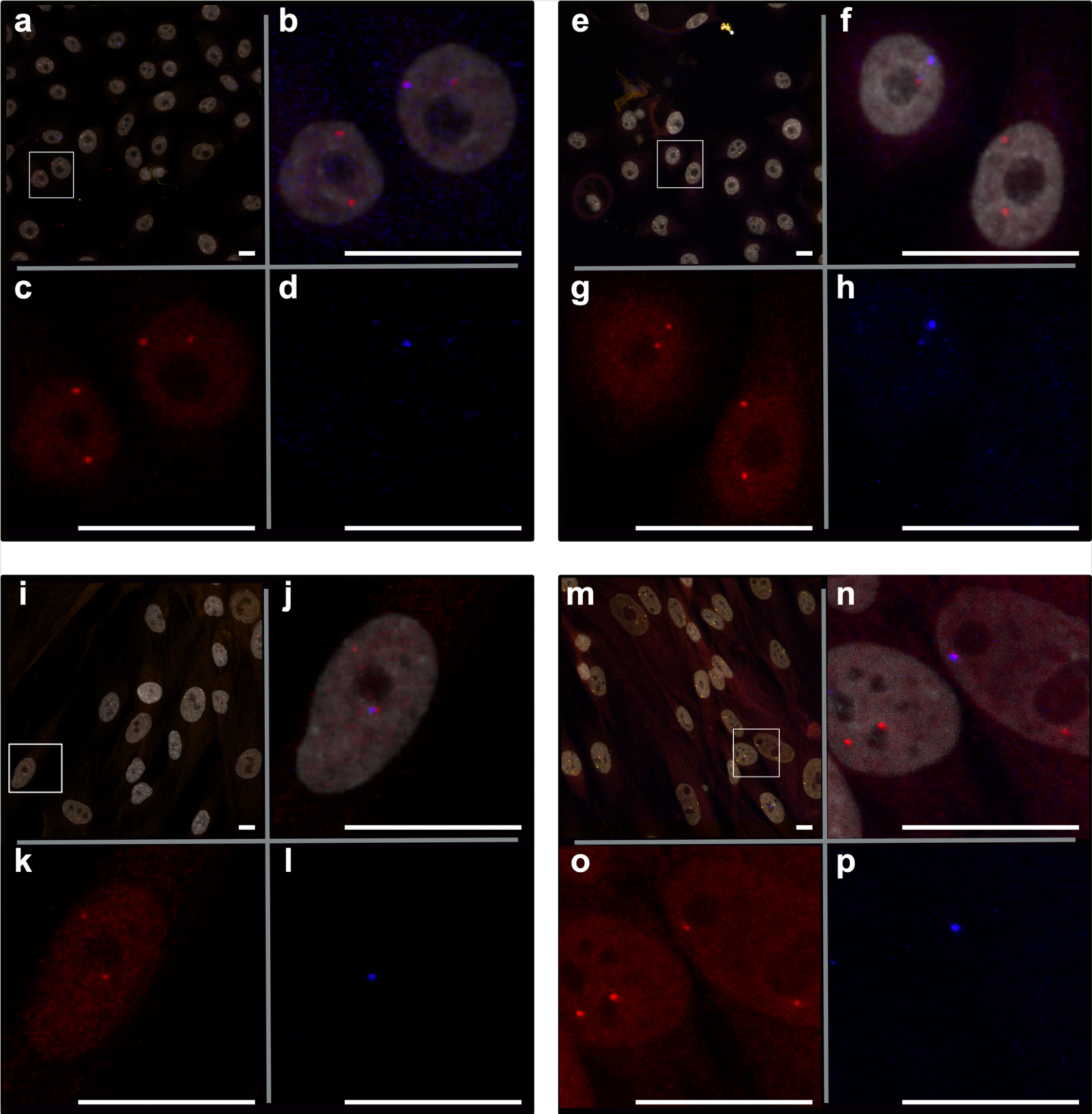
Representative images of simultaneous and sequential MYC DNA/RNA HiFISH in HBEC and HFF cells. Red: DNA, blue: nascent mRNA, grey: DAPI-stained nucleus. (a-d) simultaneous FISH in HBEC. (e-h) sequential FISH in HBEC. (i-l) simultaneous FISH in HFF. (m-p) sequential FISH in HFF. Scale bars: 20 µm

**Supplementary Figure 3:**
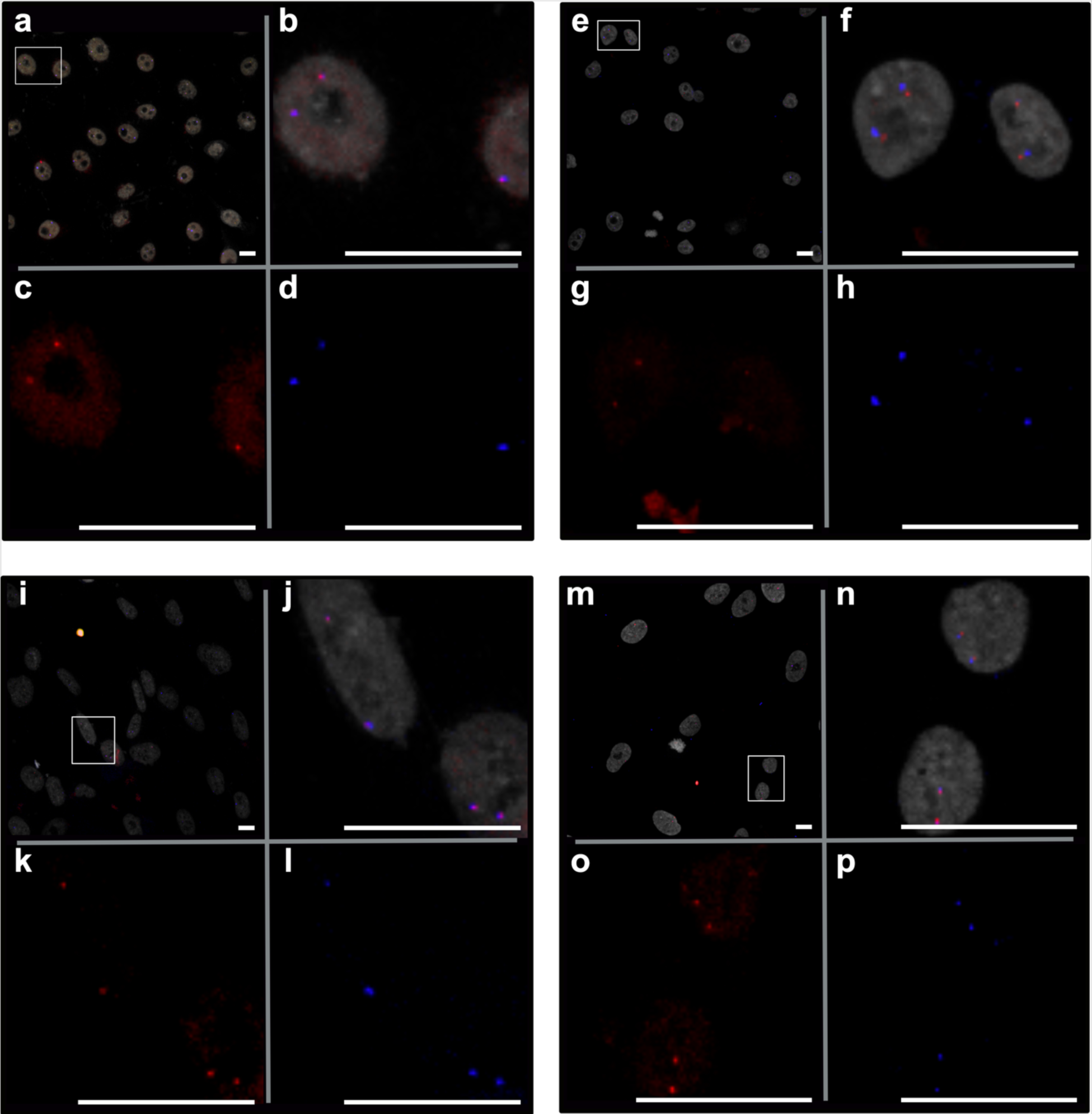
Representative images of simultaneous and sequential EGFR DNA/RNA HiFISH in HBEC and HFF cells. Red: DNA, blue: nascent mRNA, grey: DAPI-stained nucleus. (a-d) simultaneous FISH in HBEC. (e-h) sequential FISH in HBEC. (i-l) simultaneous FISH in HFF. (m-p) sequential FISH in HFF. Scale bars: 20 µm

**Supplementary Table 1.** Summary of radial distance means, standard deviations, and KS test results comparing active to inactive alleles across all experimental repeats.

| Condition |  |  |  |  | Radial distance |  |  | Two-sided Kolmogorov–Smirnov (KS) Test |  |  |
| --- | --- | --- | --- | --- | --- | --- | --- | --- | --- | --- |
| Gene | Cell-line | Protocol | Repeat | Activity | N Allele | Mean | SD | N Allele | p value | D Statistics |
| MYC | HBEC | Simultaneous | 1 | Active | 1179 | 0.7 | 0.2 | 1914.00 | 0.37 | 0.04 |
|  |  |  |  | Inactive | 735 | 0.7 | 0.2 |  |  |  |
| MYC | HBEC | Simultaneous | 2 | Active | 125 | 0.6 | 0.2 | 234.00 | 0.33 | 0.12 |
|  |  |  |  | Inactive | 109 | 0.7 | 0.2 |  |  |  |
| MYC | HBEC | Simultaneous | 3 | Active | 1629 | 0.7 | 0.2 | 2978.00 | 0.34 | 0.03 |
|  |  |  |  | Inactive | 1349 | 0.7 | 0.2 |  |  |  |
| MYC | HBEC | Sequential | 1 | Active | 781 | 0.7 | 0.2 | 1380.00 | 0.22 | 0.06 |
|  |  |  |  | Inactive | 599 | 0.7 | 0.2 |  |  |  |
| MYC | HBEC | Sequential | 2 | Active | 154 | 0.7 | 0.2 | 344.00 | 0.42 | 0.10 |
|  |  |  |  | Inactive | 190 | 0.8 | 0.2 |  |  |  |
| MYC | HFF | Simultaneous | 1 | Active | 246 | 0.7 | 0.3 | 490.00 | 0.86 | 0.05 |
|  |  |  |  | Inactive | 244 | 0.7 | 0.2 |  |  |  |
| MYC | HFF | Simultaneous | 2 | Active | 258 | 0.7 | 0.2 | 652.00 | 0.01 | 0.14 |
|  |  |  |  | Inactive | 394 | 0.7 | 0.2 |  |  |  |
| MYC | HFF | Sequential | 1 | Active | 159 | 0.7 | 0.2 | 790.00 | 0.13 | 0.10 |
|  |  |  |  | Inactive | 631 | 0.7 | 0.2 |  |  |  |
| MYC | HFF | Sequential | 2 | Active | 73 | 0.7 | 0.2 | 150.00 | 0.28 | 0.16 |
|  |  |  |  | Inactive | 77 | 0.8 | 0.2 |  |  |  |
| EGFR | HBEC | Simultaneous | 1 | Active | 3545 | 0.6 | 0.2 | 4614.00 | 0.40 | 0.03 |
|  |  |  |  | Inactive | 1069 | 0.6 | 0.2 |  |  |  |
| EGFR | HBEC | Simultaneous | 2 | Active | 7891 | 0.6 | 0.2 | 9776.00 | 0.00 | 0.05 |
|  |  |  |  | Inactive | 1885 | 0.6 | 0.2 |  |  |  |
| EGFR | HBEC | Simultaneous | 3 | Active | 7623 | 0.6 | 0.2 | 9108.00 | 0.00 | 0.05 |
|  |  |  |  | Inactive | 1485 | 0.6 | 0.2 |  |  |  |
| EGFR | HBEC | Sequential | 1 | Active | 326 | 0.6 | 0.2 | 504.00 | 0.39 | 0.08 |
|  |  |  |  | Inactive | 178 | 0.6 | 0.2 |  |  |  |
| EGFR | HBEC | Sequential | 2 | Active | 116 | 0.6 | 0.3 | 166.00 | 0.65 | 0.12 |
|  |  |  |  | Inactive | 50 | 0.6 | 0.3 |  |  |  |
| EGFR | HFF | Simultaneous | 1 | Active | 2554 | 0.6 | 0.2 | 3458.00 | 0.74 | 0.03 |
|  |  |  |  | Inactive | 904 | 0.6 | 0.2 |  |  |  |
| EGFR | HFF | Simultaneous | 2 | Active | 1184 | 0.6 | 0.3 | 1902.00 | 0.04 | 0.07 |
|  |  |  |  | Inactive | 718 | 0.6 | 0.3 |  |  |  |
| EGFR | HFF | Simultaneous | 3 | Active | 2159 | 0.6 | 0.2 | 2748.00 | 0.14 | 0.05 |
|  |  |  |  | Inactive | 589 | 0.6 | 0.2 |  |  |  |
| EGFR | HFF | Sequential | 1 | Active | 208 | 0.6 | 0.2 | 368.00 | 0.05 | 0.14 |
|  |  |  |  | Inactive | 160 | 0.7 | 0.2 |  |  |  |

Blue font color rows indicate data corresponding to Fig. 4

**Supplementary Table 2.** List of change in radial distance mean and KS test comparing simultaneous vs. sequential FISH.

| Condition |  |  |  | Radial distance | Two-sided Kolmogorov–Smirnov (KS) Test |  |  |
| --- | --- | --- | --- | --- | --- | --- | --- |
| Gene | Cell-line | Activity | Protocol | $\Delta$ mean | N allele | P value | D statistics |
| MYC | HBEC | Active | Simultaneous | 0.06 | 1960 | 0.00 | 0.23 |
|  |  |  | Sequential |  |  |  |  |
| MYC | HBEC | Inactive | Simultaneous | 0.06 | 1334 | 0.00 | 0.25 |
|  |  |  | Sequential |  |  |  |  |
| MYC | HFF | Active | Simultaneous | 0.06 | 1186 | 0.02 | 0.11 |
|  |  |  | Sequential |  |  |  |  |
| MYC | HFF | Inactive | Simultaneous | 0.04 | 1474 | 0.00 | 0.14 |
|  |  |  | Sequential |  |  |  |  |
| EGFR | HBEC | Active | Simultaneous | 0.03 | 3871 | 0.16 | 0.07 |
|  |  |  | Sequential |  |  |  |  |
| EGFR | HBEC | Inactive | Simultaneous | 0.04 | 1247 | 0.05 | 0.11 |
|  |  |  | Sequential |  |  |  |  |
| EGFR | HFF | Active | Simultaneous | 0.04 | 2762 | 0.06 | 0.10 |
|  |  |  | Sequential |  |  |  |  |
| EGFR | HFF | Inactive | Simultaneous | 0.09 | 1064 | 0.00 | 0.17 |
|  |  |  | Sequential |  |  |  |  |

